# Au_Sus: A tiered consensus census of human autophagy genes

**DOI:** 10.64898/2026.05.13.724962

**Authors:** María Guerra-Andrés, Alejandra Piedra-Macías, Ignacio García-López, Pablo Jiménez-García, Guillermo Mariño, Álvaro F. Fernández

**Affiliations:** Departamento de Bioquímica y Biología Molecular, Universidad de Oviedo, Spain; Instituto Universitario de Oncología del Principado de Asturias (IUOPA), Spain; Instituto de Investigación Sanitaria del Principado de Asturias (ISPA), Spain; Departamento de Biología Funcional, Universidad de Oviedo, Spain

**Keywords:** Autophagy, ATG genes, database, consensus, cancer signatures

## Abstract

Autophagy is a critical cellular process, yet its genomic definition remains inconsistent across digital repositories. This lack of standardisation hinders reproducibility in high-throughput studies and clinical research. Here, we present Au_Sus, a high-confidence human autophagy census established through a frequency-based majority consensus of seven primary databases and literature sources. After rigorous manual curation and nomenclature standardisation, we defined a tiered framework: Maxim_Au (2,581 genes), Au_Sus (the 201-gene core consensus), and Minim_Au (77 universal genes). Functional enrichment and protein-protein interaction analysis confirm that Au_Sus captures a highly integrated and purified autophagic machinery, with significant associations to neurodegeneration and oncology. Furthermore, an analysis of nearly 100 published cancer gene signatures revealed profound functional dilution, with 60% of signature genes absent from our consensus. These findings suggest that many of these models incorporate peripheral stress markers rather than core autophagic effectors. Hence, Au_Sus (freely accessible at <u>ausis.uniovi.es</u>) provides a reliable, ready-to-use benchmark to standardise the study of autophagy in health and disease.

## Introduction

Autophagy is a fundamental eukaryotic pathway characterised by the delivery of cytoplasmic components to the lysosome for proteolytic degradation and recycling (Galluzzi et al., 2017). Since its initial morphological description in the 1960s, our understanding of autophagy has transitioned from a basic “waste disposal” mechanism to its current status as an essential orchestrator of cellular homeostasis (Kroemer et al., 2010). Accordingly, autophagy research has become a central focus of molecular and cell biology during the last few decades, with an increasing number of studies being published every year thanks to the continuous discovery of autophagy-related (ATG) genes (Yamamoto et al., 2023). This intensive global effort has provided detailed insights into the molecular machinery of autophagy across diverse taxa, from budding yeast to complex multicellular organisms, revealing a highly conserved yet intricately regulated process (Zhang et al., 2021). The physiological significance of autophagy is perhaps best illustrated by the consequences of its failure. A robust body of literature now links autophagic dysfunction to a spectrum of human pathologies, including neurodegenerative diseases, metabolic syndromes, immune disorders, and cancer (Klionsky, Petroni, et al., 2021; Levine & Kroemer, 2019). Consequently, the modulation of the autophagic flux has emerged as a primary therapeutic target. However, as our knowledge expands, the autophagy landscape becomes increasingly complex, defined by context-specific effectors and intricate signalling cross-talk (Füllgrabe et al., 2016). To navigate this complexity, the scientific community has turned to digital resources and bioinformatic databases. Unfortunately, while these tools are indispensable for high-throughput “omics” analysis and systems biology, they have introduced new challenges regarding data reliability, consistency, and standardisation.

There are already several autophagy databases available to the scientific community (K. Chen et al., 2020; Csabai et al., 2024; Deng et al., 2018; Fu et al., 2022; Homma et al., 2011; Moussay et al., 2011; Nanduri et al., 2019; N. N. Wang et al., 2018; W. Wang et al., 2018; T. Xie et al., 2016). Each of these useful resources covers different types of information, compiling knowledge about specific organisms, regulatory mechanisms, protein-protein interactions, or chemical modulators. However, not all these lists are up to date, and they frequently include typographical errors or gene nomenclature inaccuracies. More importantly, their size can differ significantly, which means that important autophagy genes may be missing in limited databases, while extensive collections may include effectors that are only necessary in restricted pathophysiological contexts. Thus, depending on the database consulted, studies based on a single list of autophagy genes may yield divergent outcomes, contributing to the reproducibility crisis.

Here, we compare existing resources to establish a high-confidence census of essential human autophagy effectors by curating and combining the genes most consistently included across primary databases. Our goal is to provide a “consensus” list of the most relevant human autophagy protein-coding genes (Au_Sus), offering a convenient, reliable, and easy-to-use resource to advance the study of autophagy in health and disease.

## Results

### Autophagy databases show large discrepancies

To establish a comprehensive list of human autophagy effectors, we first systematically compiled available digital resources. While over ten autophagy-related databases have been reported, several of them are currently offline. Furthermore, as detailed in the Methods section, we excluded repositories that were either too restricted in scale (fewer than 150 genes) or lacked essential components of the core autophagy machinery (e.g., *ATG7*). Ultimately, seven primary sources were selected: five specialised databases (ADb, AMTDB, ANet, HADb and HAMdb) (Csabai et al., 2024; Fu et al., 2022; Homma et al., 2011; Moussay et al., 2011; N. N. Wang et al., 2018) and two literature-derived collections, hereafter referred to as *Guidelines* (Klionsky, Abdel-Aziz, et al., 2021) and *Toolbox* (Bordi et al., 2021) (Table S1).

Initial aggregation of these resources revealed significant data inconsistencies, including typographical errors, redundant entries, the presence of non-human genes, and the inclusion of protein complexes (such as AMPK or NF-kB) as single effectors rather than individual subunit-encoding genes. Consequently, we performed extensive manual curation using the HGNC multi-symbol checker, GeneCards, UniProt, and primary literature to standardise gene nomenclature and ensure all symbols reflected current genomic standards (Figure 1A). Following curation, a comparative analysis of the seven lists revealed striking discrepancies in both scale and composition. While five of the sources maintain a relatively consistent size (ranging from 604 entries in the Toolbox to 783 in HAMdb), HADb and ANet represent significant outliers (Figure 1B). Notably, HADb, which is the most frequently cited autophagy database, contains only 222 genes, while the specific subsets extracted from ANet (“Autophagy core” + “Direct regulators”) extend to 1,228 effectors.

**Figure 1.**
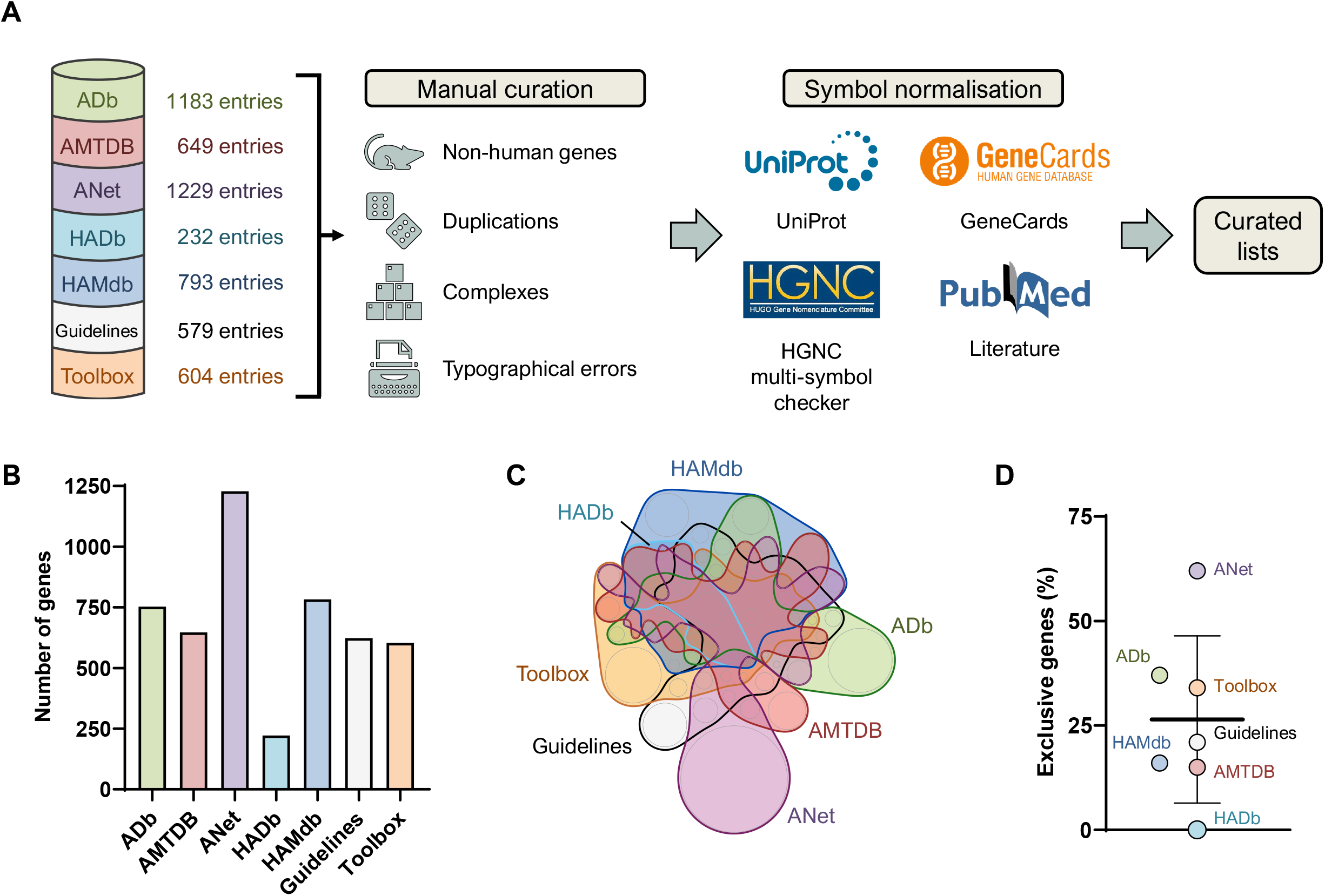
Analysis of the heterogeneity of autophagy databases. A) Overview of the workflow used for the curation of five autophagy databases and two reference articles. Manual curation was first conducted to identify and correct errors. Then, all symbols were revised and updated in accordance with the current consensus nomenclature. B) Total number of unique protein-coding genes identified in each resource following curation and symbol updating. C) Venn diagram illustrating the intersection of gene sets across all analysed resources, highlighting the significant lack of inter-database consensus. D) Representation of the percentage of genes that are unique to each specific database (Mean ± SD).

Despite these size variations, we hypothesised that the databases could still share a common core set of genes. However, divergence extends beyond list length, as shown by Figure 1C. The databases exhibit a profound lack of consensus, with nearly all resources housing large sets of genes that do not appear in the other repositories. On average, 26% of the genes in each repository are unique to that specific database (Figure 1D, Table S1). More importantly, when a functional enrichment analysis is performed on these exclusive genes, autophagy-related pathways are absent (Table S2). This suggests that these unique effectors may only be required for autophagy in highly restrictive or niche pathophysiological contexts. Pairwise comparisons further illustrate this systemic lack of overlap (Figure S1), highlighting the absence of a unified definition for human autophagy effectors. Collectively, these findings underscore a significant lack of standardisation across the autophagy repositories and demonstrate the urgent need for a curated, high-confidence consensus list to ensure the reproducibility of future studies.

### Establishment of a consensus list of human autophagy effectors

The significant lack of consensus across existing repositories underscored the need for a reliable census of key human autophagy effectors. To address this, we first integrated the entries from all seven curated resources into a single, exhaustive list. After removing redundant entries and ensuring nomenclature standardisation, this cumulative list comprised 2,581 unique protein-coding genes, which we designated as “Maxim_Au” (Figure 2A, Table S3). While Maxim_Au serves as a comprehensive catalogue of the broader autophagy landscape, its expansive nature inevitably includes effectors with putative, peripheral, transient, or highly context-specific roles in the process. Consequently, such an extensive list can dilute the functional specificity of the collection, rendering it less optimal for studies that require a focused set of primary autophagy effectors.

**Figure 2.**
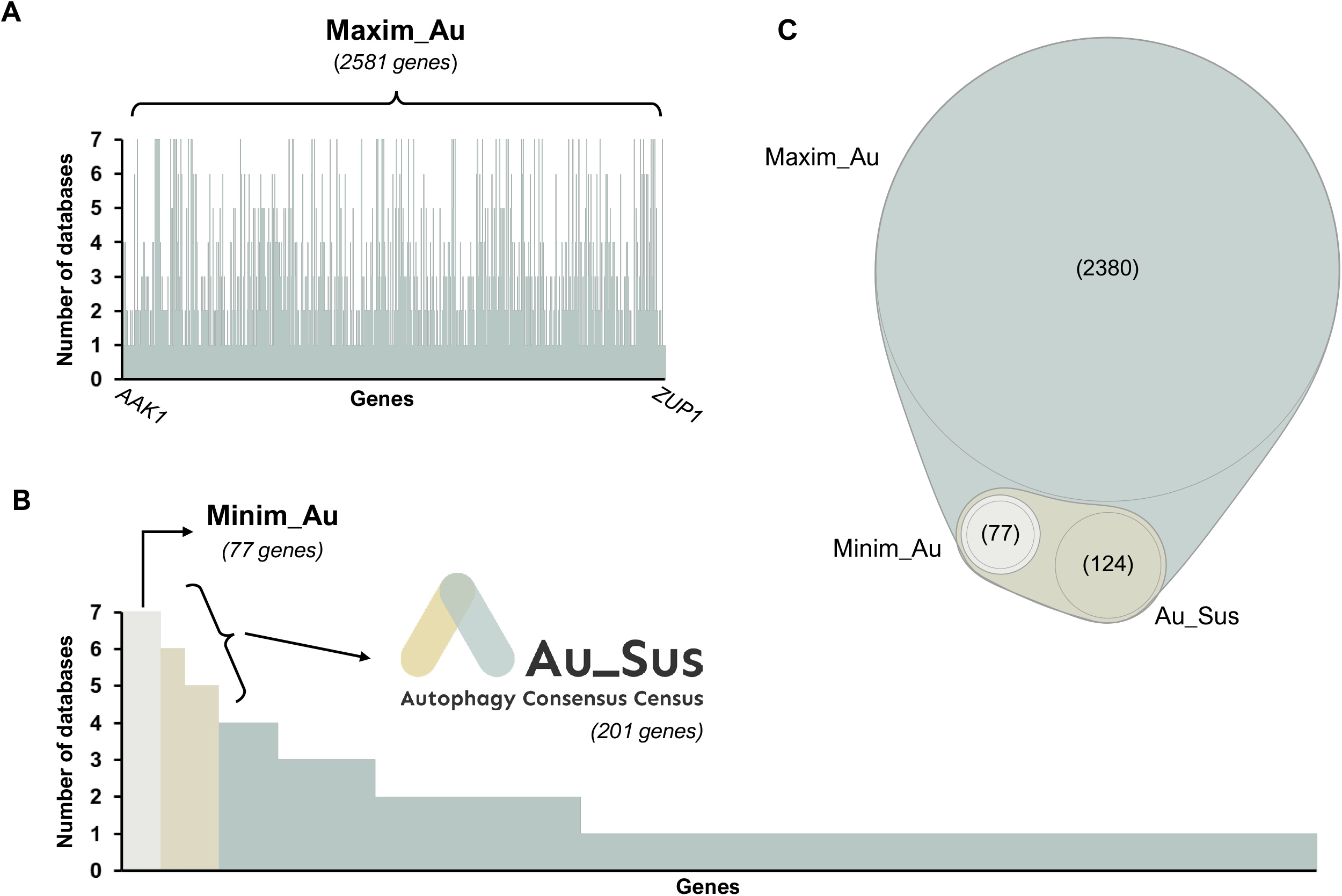
Establishment of the tiered consensus framework. A) Alphabetical distribution of the total non-redundant protein-coding genes (Maxim_Au) identified across the combined curated repositories. B) Distribution of genes based on their prevalence across the seven source databases. Genes detected in all databases were grouped into Minim_Au, whereas genes found in five or more constitute Au_Sus. C) Venn diagram illustrating the hierarchical relationship between the three proposed gene sets, offering varying levels of stringency for autophagy research.

To refine this first census, we implemented a frequency-based selection strategy (Figure 2B). We stratified each gene according to its prevalence across the seven analysed resources, hypothesising that a greater degree of overlap among databases indicates higher functional relevance and experimental validation. We then defined our primary consensus list as the subset of genes present in a robust majority (specifically, five or more) of the seven databases. This stringent threshold yielded a refined list termed “Au_Sus” (“<u>Au</u>tophagy consen<u>sus</u>”), which comprises 201 unique effectors (Table S3). By requiring a majority consensus, Au_Sus effectively eliminates inconsistent “orphan” entries while retaining the essential machinery and primary regulators of the pathway.

Finally, to provide an even more conservative resource for studies requiring absolute stringency, we identified the genes consistently present in all seven databases and defined a restrictive list of 77 effectors, termed “Minim_Au” (Figure 2B, Table S3). While Minim_Au may omit certain important regulators that have yet to be adopted across all platforms, it offers a high-purity baseline of the most universally recognised autophagic components. Together, this tiered system, ranging from the all-inclusive Maxim_Au to the high-confidence Au_Sus and the stringent Minim_Au (Figure 2C), provides a flexible and standardised toolkit for the autophagy research community, allowing investigators to select the most appropriate gene set for their specific experimental or bioinformatic context.

### Au_Sus defines a high-purity autophagy census with clinical relevance

To evaluate the representativeness of Au_Sus, we compared our consolidated list with the source databases. As shown in Figure 3A, the census displays high cross-database validation, with 86% of its effectors, on average, appearing across the seven original repositories. Notably, HADb shows an overlap of only 70% with our consensus list. In contrast, other databases such as AMTDB, HAMDb, and the Guidelines include more than 90% of the genes identified in Au_Sus. Furthermore, pairwise comparisons using Venn diagrams visually demonstrate that Au_Sus effectively overlaps with the original databases, integrating the consensus from diverse resources while filtering out the exclusive genes that lack broader support across the literature (Figure 3B).

**Figure 3.**
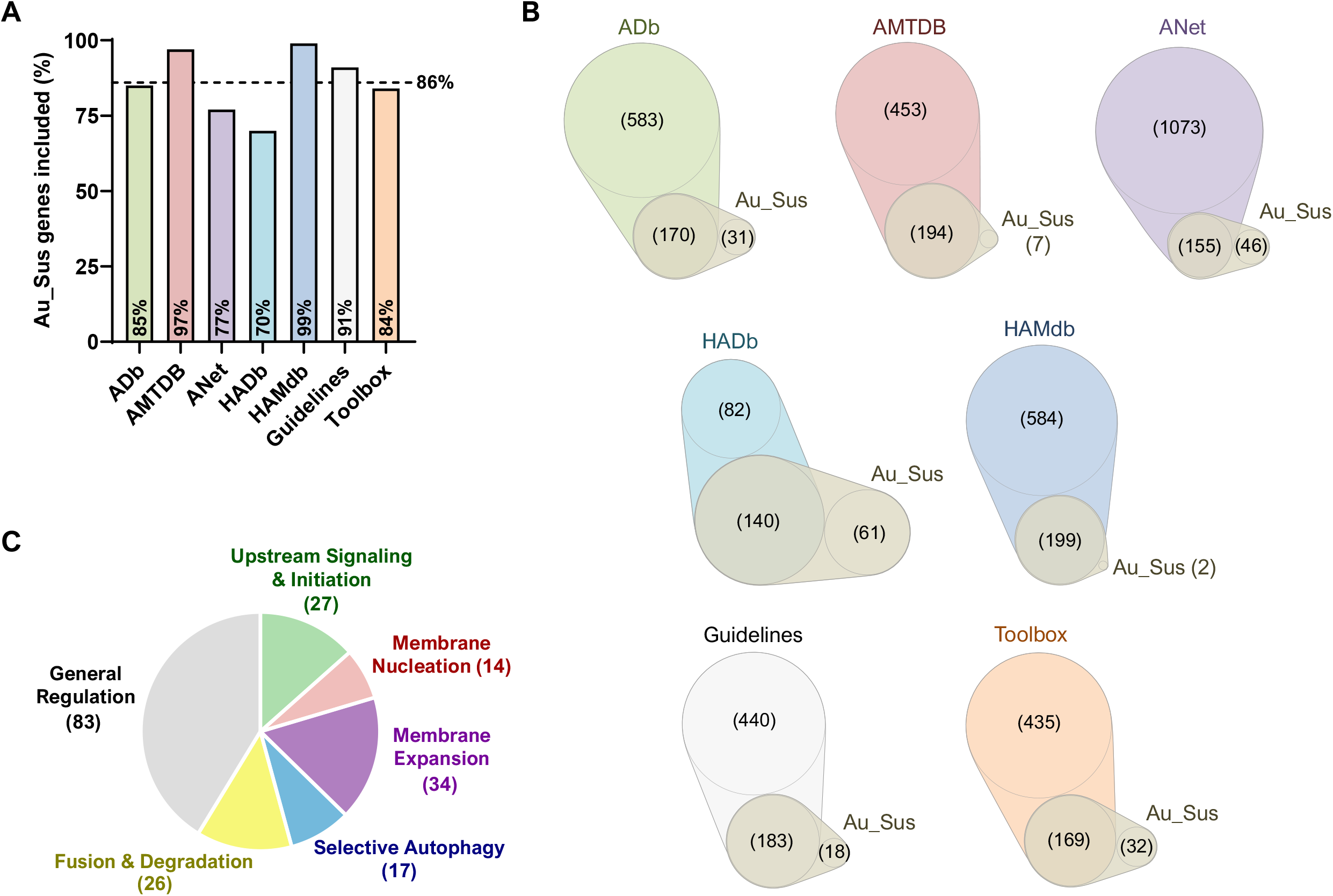
Validation of the Au_Sus census. A) Bar graph showing the percentage of Au_Sus genes represented within each original source database. The dashed horizontal line indicates the average coverage. B) Representative pairwise Venn diagrams illustrating the intersection between the Au_Sus consensus and the individual curated repositories used in this study. C) Distribution of Au_Sus effectors categorised by their canonical roles in the autophagic process, demonstrating a balanced representation of both structural machinery and regulatory elements.

To confirm that our consensus-based approach successfully isolated the core autophagy machinery from the noise of individual databases, we performed a functional enrichment analysis on the 201 genes of Au_Sus. This test shows that Au_Sus is overwhelmingly enriched in autophagy-related processes (such as “macroautophagy” or “regulation of autophagy”), but also in other terms related to cellular homeostasis and stress response (Table S4). To further characterise the molecular landscape of Au_Sus, we mapped the 201 consensus genes to the canonical stages of the autophagic process, revealing a well-balanced representation of the entire pathway (Figure 3C). The physical stages of autophagosome biogenesis (comprising its initiation, nucleation and expansion) account for roughly 37% of the census, encompassing the core ATG machinery and the early signalling complexes. But Au_Sus also provides a robust catalogue of the later stages of the route, including the selection of the cargo, as well as its final degradation. Notably, the largest functional block consists of general regulators, including transcription factors, kinases, and stress-response elements that integrate autophagy with other critical cellular processes. The distribution of these functional modules underscores that Au_Sus is not merely a list of structural components, but a comprehensive map of the autophagic system and its regulatory checkpoints. Next, to assess the functional coordination among the consensus genes, we mapped the 201 effectors of Au_Sus into a protein-protein interaction (PPI) network using the STRING database. The resulting network (Figure S2) exhibits an extraordinary level of interconnectivity, indicating that these effectors constitute a highly coordinated functional unit rather than a stochastic assembly of proteins. Its dense and modular architecture demonstrates a clear clustering effect, where genes involved in the same functional stages of autophagy are positioned in close proximity, reflecting their coordinated roles in the process. These topological features provide strong bioinformatic evidence that Au_Sus represents a functionally unified biological system rather than a random collection of genes. Finally, to evaluate the clinical relevance of our consensus list, we performed a disease association analysis using the Enrichr platform across three primary repositories: DisGeNET, OMIM, and ClinVar (Table S5-7). Our results demonstrate that Au_Sus is composed of genes with profound significance in human pathology, showing an extraordinary degree of enrichment for both neurodegenerative and neoplastic disorders. In the DisGeNET analysis (Table S5), Alzheimer’s disease emerged as the most significant association, alongside a broad spectrum of neurodegenerative disorders and various malignancies, including osteosarcoma and glioblastoma. This strong link to neurodegeneration is further supported by the OMIM hits (Table S6), which highlight characterised genetic associations with Parkinson’s disease and dementia, as well as a variety of solid tumours such as colorectal and pancreatic cancers. Lastly, the ClinVar analysis provides a direct link to pathogenicity (Table S7), revealing that Au_Sus is highly enriched in genes associated with hereditary cancer-predisposing syndromes and specific gynaecological and gastrointestinal malignancies. Collectively, these data demonstrate that our consensus approach has identified a set of essential genes that are not only central to the autophagic mechanism but also critical drivers of prevalent human diseases. This positions Au_Sus as a high-value toolkit for identifying novel therapeutic targets and biomarkers.

### Au_Sus reveals discrepancies in published cancer autophagic signatures

Given the prominence of oncology-related terms in our clinical enrichment analysis, we sought to evaluate the potential of Au_Sus as a benchmark for clinical research on autophagy and cancer. Specifically, we investigated whether the growing number of published autophagic gene signatures aligns with the high-confidence consensus established by Au_Sus. To address this, we compiled a dataset of nearly 100 published studies describing multigene autophagic signatures in cancer (Table S6). We first analysed the source for the genes used in these signatures and found that most studies relied on HADb (which omits 30% of the Au_Sus genes), while MSigDb (a repository we excluded during our curation phase for lacking essential autophagic machinery) emerged as the second most-used database (Figure 4A). This systemic reliance on potentially incomplete or inaccurate resources suggests that a significant portion of cancer research is built upon a precarious bioinformatic foundation. This concern was confirmed by a comparative overlap analysis between the total pool of genes identified across these signatures (311 unique effectors) and our consensus list. The results revealed a profound discrepancy in gene composition, as 62,7% of the genes included in published cancer signatures are entirely absent from Au_Sus (Figure 4B). Crucially, only four of the ten genes most included in all the signatures are part of Au_Sus (with none of them appearing in Minim_Au) (Figure 4C). Moreover, when we performed functional enrichment analysis on them, autophagy-related processes were absent, with the top hits being related to signalling regulation and cell death (mostly apoptosis) (Figure 4D, Table S7). This implies that these models incorporate genes that lack a broad consensus as primary autophagic effectors, likely capturing peripheral biomarkers of cellular stress or tumour progression rather than the autophagic process itself. These findings highlight a systemic issue in clinical bioinformatics: the “autophagic status” of a tumour is often determined by genes that may not be central to the pathway. By providing a curated, high-confidence benchmark, Au_Sus offers a solution to mitigate this functional dilution, ensuring that future cancer signatures are biologically accurate and reproducible.

**Figure 4.**
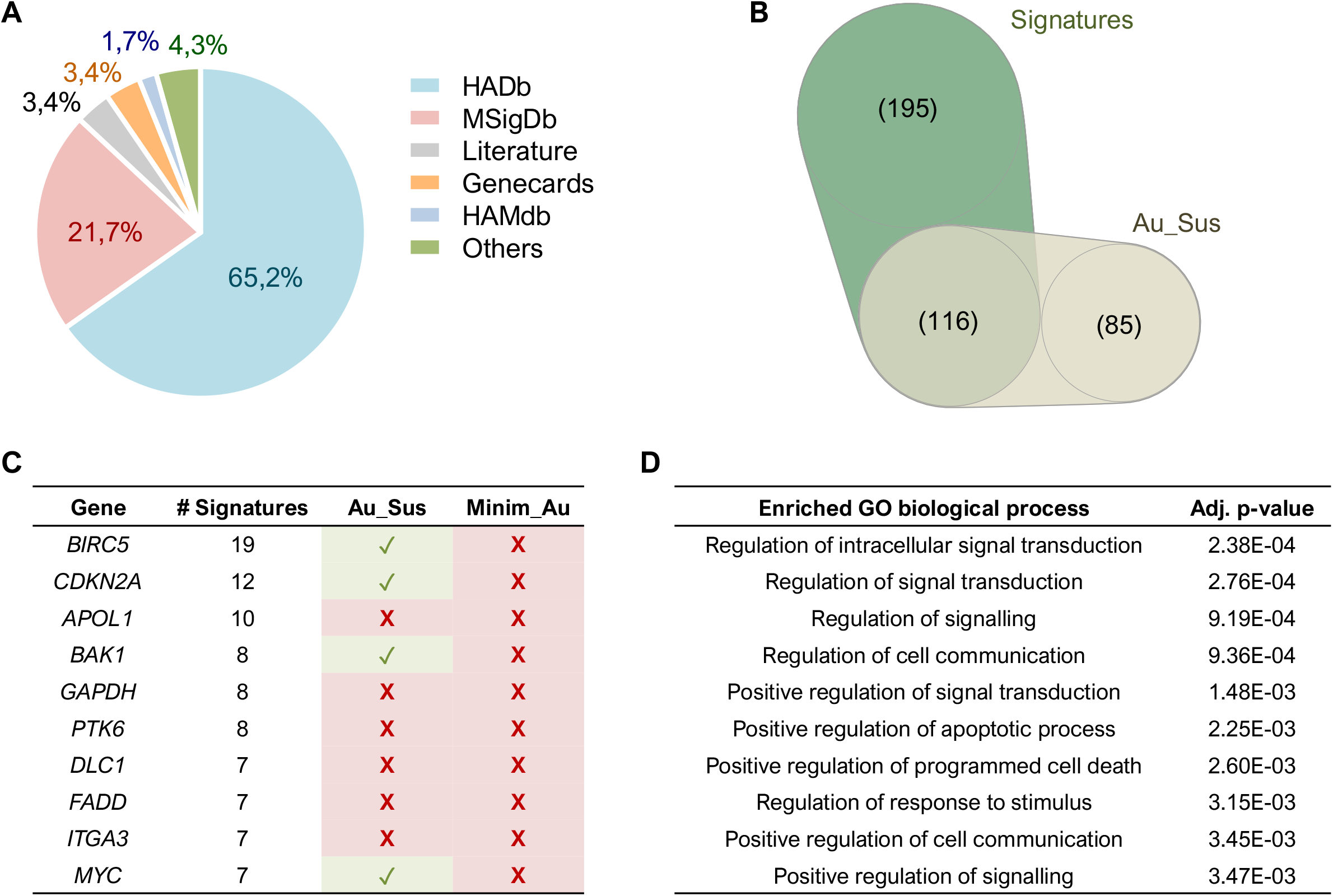
Au_Sus reveals functional dilution in published cancer autophagic signatures. A) Distribution of source databases used in the construction of nearly 100 published cancer autophagic signatures. B) Venn diagram showing the overlap between the total pool of signature-derived genes and the Au_Sus census. C) Most frequent genes in published cancer autophagic signatures and their presence or absence in the Au_Sus list. None of them is part of Minim_Au. D) Pathway enrichment analysis on the most frequent genes included in the signatures of the compiled publications, showing the absence of the autophagic process.

## Discussion

The autophagy community is currently supported by several databases that function as deep-dive encyclopaedias. Resources such as HADb, HAMdb, and ANet provide invaluable specialised information, including protein-protein interactions, post-translational modifications, and chemical modulators. Therefore, these platforms remain essential for researchers investigating the specific molecular details of autophagy proteins. However, the sheer volume of information they contain can make it difficult for researchers to quickly extract a concise, standardised list of human autophagy effectors for high-throughput “omics” studies or clinical benchmarking. Indeed, while the rapid expansion of the autophagy field has generated a substantial amount of digital data, this abundance has introduced a significant challenge regarding genomic standardisation. As our analysis demonstrates, the lack of a unified consensus among primary autophagy databases has led to a fragmented landscape where the “autophagic identity” of a gene often depends on the specific repository consulted. Au_Sus was developed to address this discrepancy, providing a tiered, frequency-based census that identifies a high-confidence core of human autophagy effectors for immediate practical application, serving as a complementary work tool along with other autophagy databases. By applying a majority-consensus threshold, we have filtered out the “noise” of context-specific or transient interactors, resulting in a census with an extraordinary degree of functional enrichment. This ensures that researchers using Au_Sus are working with a list that is both comprehensive and biologically robust.

To evaluate the real-world implications of this lack of standardisation and validate the potential of Au_Sus, we analysed nearly 100 published cancer signatures. This analysis revealed that more than 60% of their genes are absent from our consensus list, suggesting that they are likely tracking biomarkers of general cellular stress or cancer progression rather than true autophagic executioners. While these “non-consensus” genes may correlate with disease outcomes, their frequent mislabelling as primary autophagy effectors can lead to misleading conclusions in translational research. By providing a curated benchmark, Au_Sus allows for the refinement of these signatures, ensuring that clinical observations are grounded in the actual molecular machinery of the autophagic process.

We envision Au_Sus as a fundamental resource for researchers navigating the complex landscape of autophagy in health and disease. Thus, to facilitate its widespread adoption, Au_Sus (along with the Maxim_Au and Minim_Au tiers) is freely available for download at our webpage: <u>ausis.uniovi.es</u>. However, we recognise that a bioinformatic census is only as useful as its currency. Therefore, we are committed to maintaining Au_Sus as a living repository to ensure its long-term utility. We will perform periodic reviews and update the framework whenever old source databases undergo significant revisions or new ones are published, ensuring that the scientific community has access to a list that reflects the most up-to-date consensus in the field. Still, certain limitations remain. As a frequency-based model, Au_Sus may exclude genes that could be relevant in specific tissues or rare physiological conditions. Furthermore, our current focus is restricted to protein-coding genes. As the roles of non-coding RNAs and epigenetic regulators in autophagy become more clearly defined, future iterations of the census may need to expand to include these vital regulatory layers.

## Methods

### Data selection and database comparison

Human autophagic genes were retrieved from five different databases: the Human Autophagy Database (HADb, http://autophagy.lu), the Human Autophagy Modulator Database (HAMdb, http://hamdb.scbdd.com), the Autophagy Database (ADb, http://www.tanpaku.org/autophagy.html), the Autophagic Modulators in Tumours Database (AMTDB, https://amtdb.vercel.app), and the “Autophagy core” + “Direct regulators” subsets that are included in AutophagyNet (https://www.autophagynet.org/download). Autophagy effectors were also collected from two additional documents: the glossary in the *Guidelines* (a recurring publication that recapitulates our knowledge of autophagy) (Klionsky, Abdel-Aziz, et al., 2021), and the gene toolbox proposed by Bordi and collaborators (Bordi et al., 2021). Other autophagy databases (such as the Autophagy Compound Database (ACDB, http://www.acdbliulab.com), the Autophagy Compound-Target Prediction database (ACTP, http://actp.liu-lab.com), the Autophagy to Disease database (ATD, http://auto2disease.nwsuaflmz.com), the Autophagy Small Molecule Database (ASMDB, http://www.autophagysmdb.org), the Autophagy and Tumor Database (ATdb, http://www.bigzju.com/ATdb/#), the autophagy gene set at the Molecular Signature Database (MSigDb, https://www.gsea-msigdb.org/gsea/msigdb), or Reactome (https://reactome.org)) were discarded either because 1) the repository is no longer available, 2) the gene list is too short (< 150 genes), or 3) the collection lacks indispensable genes of the core autophagy machinery (like *ATG7*). Direct comparisons between databases or sets of genes were plotted with the nVenn tool (https://degradome.uniovi.es/cgi-bin/nVenn/nvenn.cgi) (Pérez-Silva et al., 2018).

### Gene nomenclature standardisation and manual curation

All collected entries underwent rigorous manual curation to ensure genomic standardisation. We restricted the census to human protein-coding genes. To resolve discrepancies in nomenclature and redundant entries, all symbols were processed through the HUGO Gene Nomenclature Committee (HGNC) multi-symbol checker (https://www.genenames.org/tools/multi-symbol-checker) (Seal et al., 2025). Ambiguous symbols or historical aliases were further cross-referenced using GeneCards (https://www.genecards.org) (Stelzer et al., 2016) and UniProt (https://www.uniprot.org) (Bateman et al., 2025) to ensure all effectors were assigned the current official symbols.

### Generation of stratified autophagy lists

The curated lists were integrated into a cumulative dataset, Maxim_Au, representing the union of all seven sources. To identify the high-confidence core of the pathway, we employed a frequency-based majority consensus strategy. Each gene was ranked by its prevalence across the seven sources. The primary consensus list, Au_Sus, was defined as the subset of genes present in five or more databases. This threshold was determined by analysing the gene distribution frequency, as a significant expansion in gene count was observed when lowering the stringency to four databases (where the number of “exclusive” genes was nearly double that of the five-database threshold). For studies requiring absolute stringency, we established Minim_Au, comprising only those genes present in all seven databases.

### Validation analyses

Functional enrichment analysis was performed using g:Profiler (GO:BP) (https://biit.cs.ut.ee/gprofiler/gost), annotating the resulting adjusted p-value (Kolberg et al., 2023). To assess molecular connectivity, the Au_Sus census was queried against the STRING database (v12.0, https://string-db.org) (Szklarczyk et al., 2023), utilising a standard medium confidence score (0.400). The network’s topological robustness was evaluated based on the PPI enrichment p-value, average node degree, and local clustering coefficient. Disease association analysis was conducted using the Enrichr platform (https://maayanlab.cloud/Enrichr/), querying the DisGeNET, OMIM, and ClinVar repositories (E. Y. Chen et al., 2013; Kuleshov et al., 2016; Z. Xie et al., 2021). To evaluate the practical application of our census, a systematic search was performed in PubMed (up to 2026) to identify the establishment of multigene autophagic signatures for cancer. Search terms included “autophagy”, “gene signature”, “prognosis” and “cancer”. Studies were included if they provided a discrete list of gene symbols used to construct a predictive model. For each study, the sources used to retrieve the initial gene set (e.g., HADb, MSigDb) were recorded. A total of 92 unique signatures were curated, resulting in a meta-pool of 311 unique gene effectors. This set of signature-derived genes was then cross-referenced against the Au_Sus census. Genes were categorised as “consensus-validated” if present in both lists, or “signature-exclusive” if absent from the consensus. Functional enrichment analysis of top signature genes was conducted to identify potential functional dilution, as previously described.

## Author Contributions

Conceptualization, Á.F.F.; data acquisition and curation, M.G.A. and Á.F.F; data analysis, M.G.A, A.P.M., I.G.L., P.J.G. and Á.F.F.; writing (original draft preparation), M.G.A. and Á.F.F.; writing (review and editing), M.G.A., A.P.M., G.M. and Á.F.F.; supervision, Á.F.F. All authors have read and agreed to the published version of the manuscript.

## Acknowledgements

We thank Dr Víctor Quesada for assistance with his nVenn tool. We are members of and thank Women in Autophagy (WIA), Sociedad Española de Bioquímica y Biología Molecular (SEBBM) and Sociedad Española de Autofagia (SEFAGIA).

## References

Bateman, A., Martin, M. J., Orchard, S., Magrane, M., Adesina, A., Ahmad, S., Bowler-Barnett, E. H., Bye-A-Jee, H., Carpentier, D., Denny, P., Fan, J., Garmiri, P., da Costa Gonzales, L. J., Hussein, A., Ignatchenko, A., Insana, G., Ishtiaq, R., Joshi, V., Jyothi, D., … Zhang, J. (2025). UniProt: the Universal Protein Knowledgebase in 2025. Nucleic Acids Research, 53(D1), D609–D617. 10.1093/NAR/GKAE1010

Bordi, M., De Cegli, R., Testa, B., Nixon, R. A., Ballabio, A., & Cecconi, F. (2021). A gene toolbox for monitoring autophagy transcription. Cell Death & Disease, 12(11), 1044. 10.1038/s41419-021-04121-9

Chen, E. Y., Tan, C. M., Kou, Y., Duan, Q., Wang, Z., Meirelles, G. V., Clark, N. R., & Ma’ayan, A. (2013). Enrichr: interactive and collaborative HTML5 gene list enrichment analysis tool. BMC Bioinformatics 2013 14:1, 14(1), 128-. 10.1186/1471-2105-14-128

Chen, K., Yang, D., Zhao, F., Wang, S., Ye, Y., Sun, W., Lu, H., Ruan, Z., Xu, J., Wang, T., Lu, G., Wang, L., Shi, Y., Zhang, H., Wu, H., Lu, W., Shen, H. M., Xia, D., & Wu, Y. (2020). Autophagy and Tumor Database: ATdb, a novel database connecting autophagy and tumor. Database, 2020, 1–11. 10.1093/database/baaa052

Csabai, L., Bohár, B., Türei, D., Prabhu, S., Földvári-Nagy, L., Madgwick, M., Fazekas, D., Módos, D., Ölbei, M., Halka, T., Poletti, M., Kornilova, P., Kadlecsik, T., Demeter, A., Szalay-Bekő, M., Kapuy, O., Lenti, K., Vellai, T., Gul, L., & Korcsmáros, T. (2024). AutophagyNet: high-resolution data source for the analysis of autophagy and its regulation. Autophagy, 20(1), 188–201. 10.1080/15548627.2023.2247737

Deng, Y., Zhu, L., Cai, H., Wang, G., & Liu, B. (2018). Autophagic compound database: A resource connecting autophagy-modulating compounds, their potential targets and relevant diseases. Cell Proliferation, 51(3). 10.1111/CPR.12403

Fu, J., Wu, L., Hu, G., Shi, Q., Wang, R., Zhu, L., Yu, H., & Fu, L. (2022). AMTDB: A comprehensive database of autophagic modulators for anti-tumor drug discovery. Frontiers in Pharmacology, 0, 3005. 10.3389/FPHAR.2022.956501

Füllgrabe, J., Ghislat, G., Cho, D. H., & Rubinsztein, D. C. (2016). Transcriptional regulation of mammalian autophagy at a glance. Journal of Cell Science, 129(16), 3059–3066. 10.1242/jcs.188920

Galluzzi, L., Baehrecke, E. H., Ballabio, A., Boya, P., Bravo-San Pedro, J. M., Cecconi, F., Choi, A. M., Chu, C. T., Codogno, P., Colombo, M. I., Cuervo, A. M., Debnath, J., Deretic, V., Dikic, I., Eskelinen, E., Fimia, G. M., Fulda, S., Gewirtz, D. A., Green, D. R., … Kroemer, G. (2017). Molecular definitions of autophagy and related processes. The EMBO Journal, 36(13), 1811–1836. 10.15252/embj.201796697

Homma, K., Suzuki, K., & Sugawara, H. (2011). The autophagy database: An all-inclusive information resource on autophagy that provides nourishment for research. Nucleic Acids Research, 39(SUPPL. 1), 986–990. 10.1093/nar/gkq995

Klionsky, D. J., Abdel-Aziz, A. K., Abdelfatah, S., Abdellatif, M., Abdoli, A., Abel, S., Abeliovich, H., Abildgaard, M. H., Abudu, Y. P., Acevedo-Arozena, A., Adamopoulos, I. E., Adeli, K., Adolph, T. E., Adornetto, A., Aflaki, E., Agam, G., Agarwal, A., Aggarwal, B. B., Agnello, M., … Tong, C. K. (2021). Guidelines for the use and interpretation of assays for monitoring autophagy (4th edition). Autophagy. 10.1080/15548627.2020.1797280

Klionsky, D. J., Petroni, G., Amaravadi, R. K., Baehrecke, E. H., Ballabio, A., Boya, P., Bravo-San Pedro, J. M., Cadwell, K., Cecconi, F., Choi, A. M. K., Choi, M. E., Chu, C. T., Codogno, P., Colombo, M. I., Cuervo, A. M., Deretic, V., Dikic, I., Elazar, Z., Eskelinen, E., … Pietrocola, F. (2021). Autophagy in major human diseases. The EMBO Journal, 40(19), EMBJ2021108863-. 10.15252/EMBJ.2021108863/FIGURES/2

Kolberg, L., Raudvere, U., Kuzmin, I., Adler, P., Vilo, J., & Peterson, H. (2023). g:Profiler— interoperable web service for functional enrichment analysis and gene identifier mapping (2023 update). Nucleic Acids Research, 51(W1), W207–W212. 10.1093/NAR/GKAD347

Kroemer, G., Mariño, G., & Levine, B. (2010). Autophagy and the Integrated Stress Response. Molecular Cell, 40(2), 280–293. 10.1016/j.molcel.2010.09.023

Kuleshov, M. V., Jones, M. R., Rouillard, A. D., Fernandez, N. F., Duan, Q., Wang, Z., Koplev, S., Jenkins, S. L., Jagodnik, K. M., Lachmann, A., McDermott, M. G., Monteiro, C. D., Gundersen, G. W., & Maayan, A. (2016). Enrichr: a comprehensive gene set enrichment analysis web server 2016 update. Nucleic Acids Research, 44(W1), W90–W97. 10.1093/NAR/GKW377

Levine, B., & Kroemer, G. (2019). Biological Functions of Autophagy Genes: A Disease Perspective. Cell, 176(1–2), 11–42. 10.1016/J.CELL.2018.09.048

Moussay, E., Kaoma, T., Baginska, J., Muller, A., Van Moer, K., Nicot, N., Nazarov, P. V., Vallar, L., Chouaib, S., Berchem, G., & Janji, B. (2011). The acquisition of resistance to TNFα in breast cancer cells is associated with constitutive activation of autophagy as revealed by a transcriptome analysis using a custom microarray. Autophagy, 7(7), 760–770. 10.4161/auto.7.7.15454

Nanduri, R., Kalra, R., Bhagyaraj, E., Chacko, A. P., Ahuja, N., Tiwari, D., Kumar, S., Jain, M., Parkesh, R., & Gupta, P. (2019). AutophagySMDB: a curated database of small molecules that modulate protein targets regulating autophagy. Autophagy, 15(7), 1280–1295. 10.1080/15548627.2019.1571717

Pérez-Silva, J. G., Araujo-Voces, M., & Quesada, V. (2018). nVenn: generalized, quasi-proportional Venn and Euler diagrams. Bioinformatics, 34(13), 2322–2324. 10.1093/BIOINFORMATICS/BTY109

Seal, R. L., Braschi, B., Gray, K., McClay, J., Tweedie, S., & Bruford, E. A. (2025). Genenames.org: the HGNC and PGNC resources in 2026. Nucleic Acids Research, 54(D1), D1098. 10.1093/NAR/GKAF1229

Stelzer, G., Rosen, N., Plaschkes, I., Zimmerman, S., Twik, M., Fishilevich, S., Iny Stein, T., Nudel, R., Lieder, I., Mazor, Y., Kaplan, S., Dahary, D., Warshawsky, D., Guan-Golan, Y., Kohn, A., Rappaport, N., Safran, M., & Lancet, D. (2016). The GeneCards suite: From gene data mining to disease genome sequence analyses. Current Protocols in Bioinformatics, 2016(1), 1.30.1–1.30.33. 10.1002/CPBI.5;JOURNAL:JOURNAL:1934340X;WGROUP:STRING:PUBLICATION

Szklarczyk, D., Kirsch, R., Koutrouli, M., Nastou, K., Mehryary, F., Hachilif, R., Gable, A. L., Fang, T., Doncheva, N. T., Pyysalo, S., Bork, P., Jensen, L. J., & Von Mering, C. (2023). The STRING database in 2023: protein–protein association networks and functional enrichment analyses for any sequenced genome of interest. Nucleic Acids Research, 51(D1), D638–D646. 10.1093/NAR/GKAC1000

Wang, N. N., Dong, J., Zhang, L., Ouyang, D., Cheng, Y., Chen, A. F., Lu, A. P., & Cao, D. S. (2018). HAMdb: a database of human autophagy modulators with specific pathway and disease information. Journal of Cheminformatics, 10(1), 1–8. 10.1186/s13321-018-0289-4

Wang, W., Zhang, P., Li, L., Chen, Z., Bai, W., Liu, G., Zhang, L., Jia, H., Li, L., Yu, Y., & Liao, M. (2018). ATD: A comprehensive bioinformatics resource for deciphering the association of autophagy and diseases. Database, 2018(2018), 1–10. 10.1093/database/bay093

Xie, T., Zhang, L., Zhang, S., Ouyang, L., Cai, H., Liu, B., Xie, T., Zhang, L., Zhang, S., Ouyang, L., Cai, H., & Liu, B. (2016). ACTP: A webserver for predicting potential targets and relevant pathways of autophagymodulating compounds. Oncotarget, 7(9), 10015–10022. 10.18632/ONCOTARGET.7015

Xie, Z., Bailey, A., Kuleshov, M. V., Clarke, D. J. B., Evangelista, J. E., Jenkins, S. L., Lachmann, A., Wojciechowicz, M. L., Kropiwnicki, E., Jagodnik, K. M., Jeon, M., & Ma’ayan, A. (2021). Gene Set Knowledge Discovery with Enrichr. Current Protocols, 1(3), e90. 10.1002/CPZ1.90

Yamamoto, H., Zhang, S., & Mizushima, N. (2023). Autophagy genes in biology and disease. Nature Reviews Genetics 2023 24:6, 24(6), 382–400. 10.1038/s41576-022-00562-w

Zhang, S., Hama, Y., & Mizushima, N. (2021). The evolution of autophagy proteins – Diversification in eukaryotes and potential ancestors in prokaryotes. Journal of Cell Science, 134(13). 10.1242/JCS.233742/270774

