## Supplementary Figures for "Au_Sus: A tiered consensus census of human autophagy genes"

### Supplementary figure legends

**Supplementary Figure 1. Pairwise gene overlap analysis across the seven curated repositories.** Venn diagrams illustrating the intersection of gene sets between all possible pairwise combinations of the seven source databases. The widespread lack of substantial overlap underscores the significant heterogeneity in how the human autophagic machinery is currently catalogued across digital resources.

**Supplementary Figure 2. Functional connectivity and modular architecture of the Au_Sus census.** Protein–protein interaction (PPI) network of the 201 effectors included in Au_Sus, generated using the STRING database. The network displays a highly integrated and non-stochastic topology, where proteins exhibit a clear clustering effect based on their coordinated biological roles. Nodes are colour-coded according to their canonical stage in the autophagic process: upstream signalling and initiation (green), nucleation (red), elongation (purple), selective autophagy (blue), fusion and degradation (yellow), and general regulation (grey). This spatial distribution demonstrates that functionally related effectors maintain close topological proximity within the consensus core.

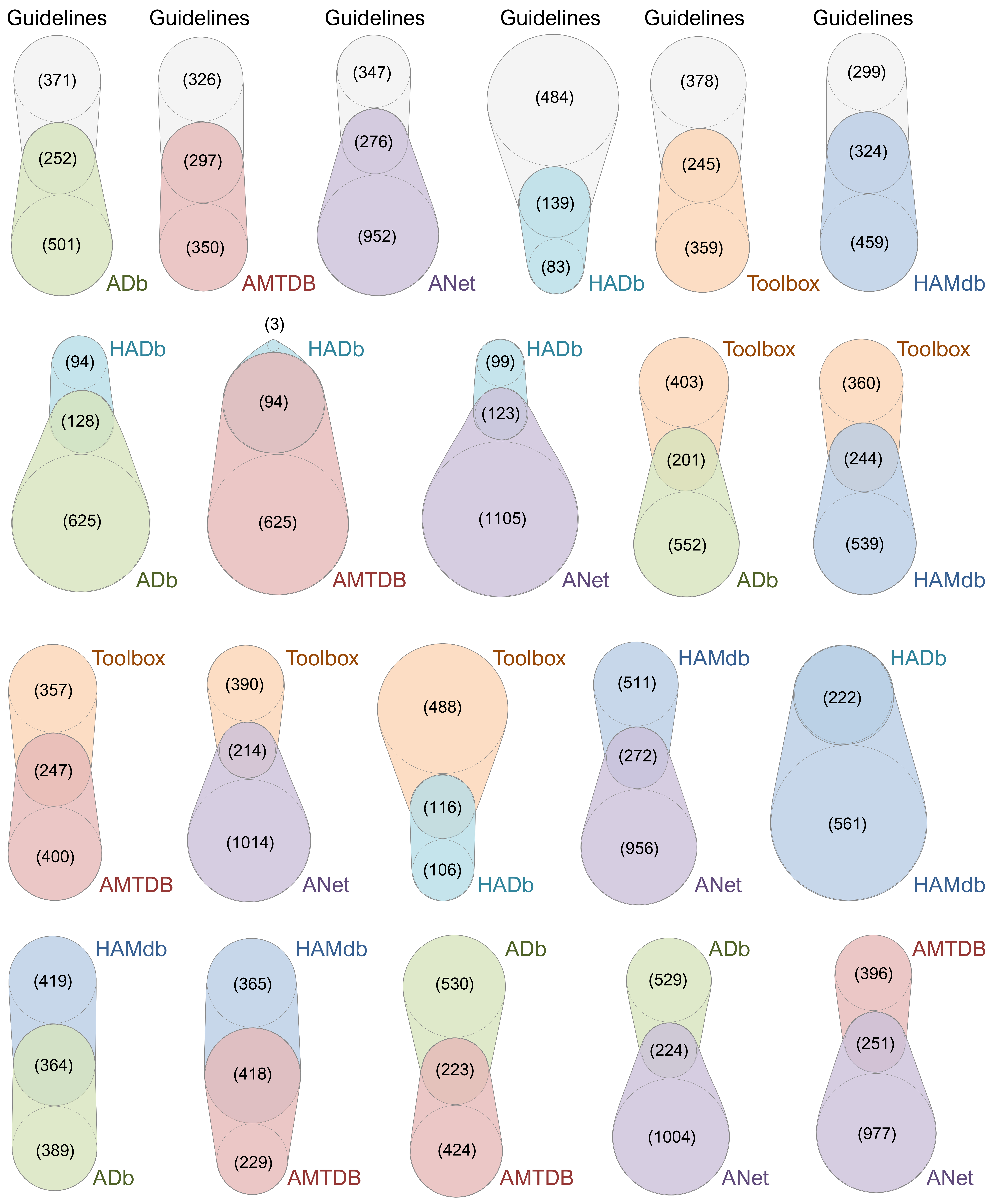
Guidelines
(371)
(252)
(501)
ADb
Guidelines
(326)
(297)
(350)
AMTDB
Guidelines
(347)
(276)
(952)
ANet
Guidelines
(484)
(139)
(83)
HADb
Guidelines
(378)
(245)
(359)
Toolbox
Guidelines
(299)
(324)
(459)
HAMdb
HADb
(94)
(128)
(625)
ADb
(3)
HADb
(94)
(625)
AMTDB
HADb
(99)
(123)
(1105)
ANet
Toolbox
(403)
(201)
(552)
ADb
Toolbox
(360)
(244)
(539)
HAMdb
Toolbox
(357)
(247)
(400)
AMTDB
Toolbox
(390)
(214)
(1014)
ANet
Toolbox
(488)
(116)
(106)
HADb
HAMdb
(511)
(272)
(956)
ANet
HADb
(222)
(561)
HAMdb
HAMdb
(419)
(364)
(389)
ADb
HAMdb
(365)
(418)
(229)
AMTDB
ADb
(530)
(223)
(424)
AMTDB
ADb
(529)
(224)
(1004)
ANet
AMTDB
(396)
(251)
(977)
ANet

**Figure S1**

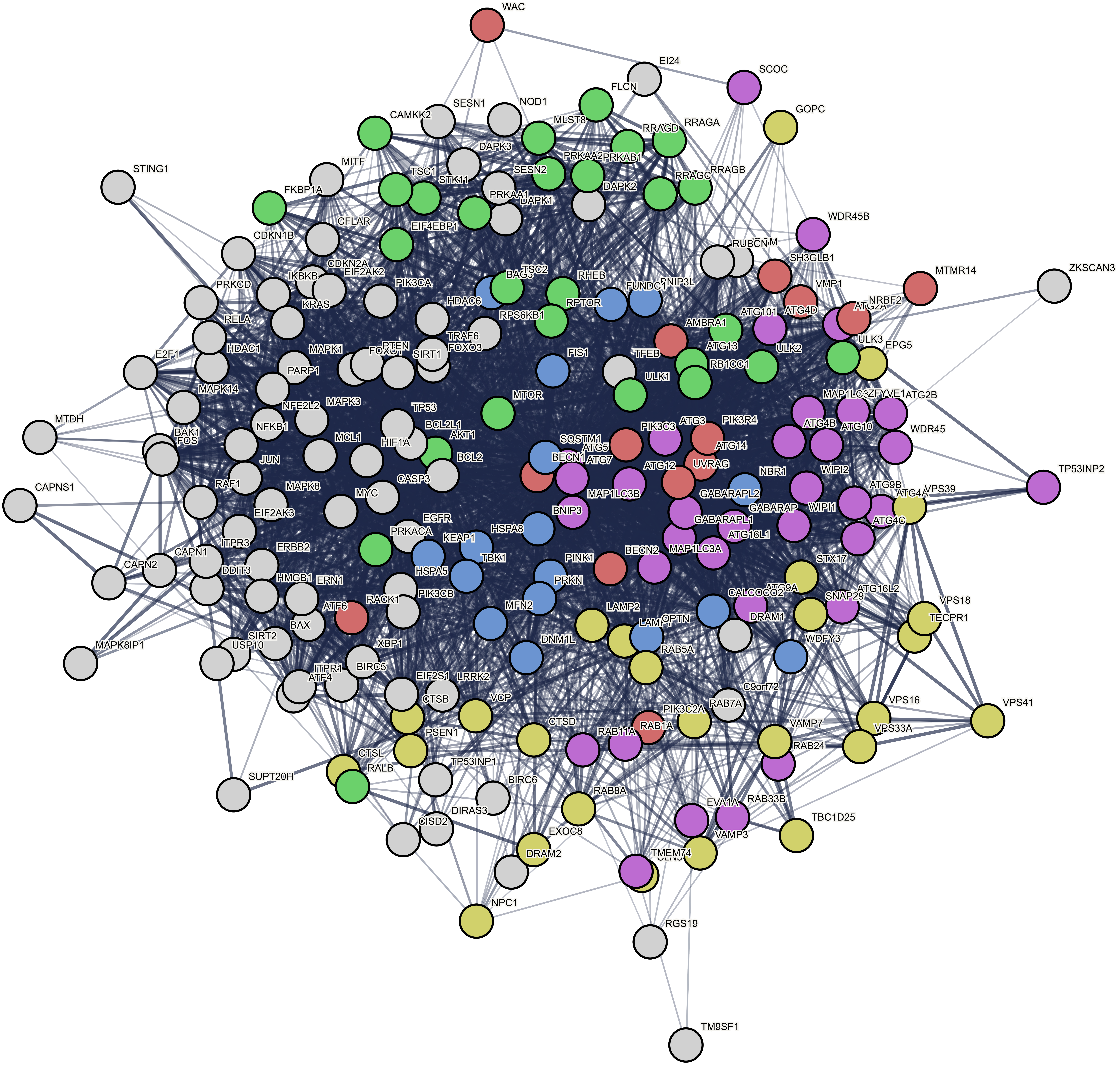


| | |
|---|---|
| Number of nodes | 201 |
| Number of edges | 4851 |
| Average node degree | 48.3 |
| Avg. local clustering coefficient | 0.643 |
| PPI enrichment p-value | <1.0E-16 |

**Figure S2**
